# Deviations from an individual’s average sociality have fitness consequences

**DOI:** 10.64898/2026.05.28.728517

**Authors:** Taylor N. Bastian, Conner S. Philson, Julien G. A. Martin, Daniel T. Blumstein

## Abstract

Social relationships are consequential and affect survival, longevity and reproductive success in many species. Prior studies have successfully identified the fitness consequences of variation in sociality relative to other individuals. However, we have yet to quantify the fitness consequences of social variation within an individual over time. By using a within-individual centering approach, we can identify the consequences of deviations from an individual’s ‘average’ sociality. This approach has not been applied to the study of animal sociality. We used 20 years of data from a population of yellow-bellied marmots (*Marmota flaviventer*) to understand the fitness consequences of an individual’s average affiliative social network position over their lifetime, and of individuals’ annual deviations from their average social position. We found that an individual’s deviation from its average sociality was very strongly associated with both winter and summer survival. Deviations from some average measures of sociality were also associated with the likelihood of reproducing, but not in a consistent direction. Lifetime average social network positions had limited associations with survival and reproduction. Interestingly, litter size results varied between males to females; neither lifetime averages of social network traits nor deviations from these averages were associated with litter size in females, but deviations in sociality were associated with larger numbers of weaned offspring in males. Future studies of the consequences of social relationships may benefit from the use of an individual-centering approach, and further work is required to understand the environmental and social drivers of this intra-individual adaptive plasticity.

**Graphical abstract:** 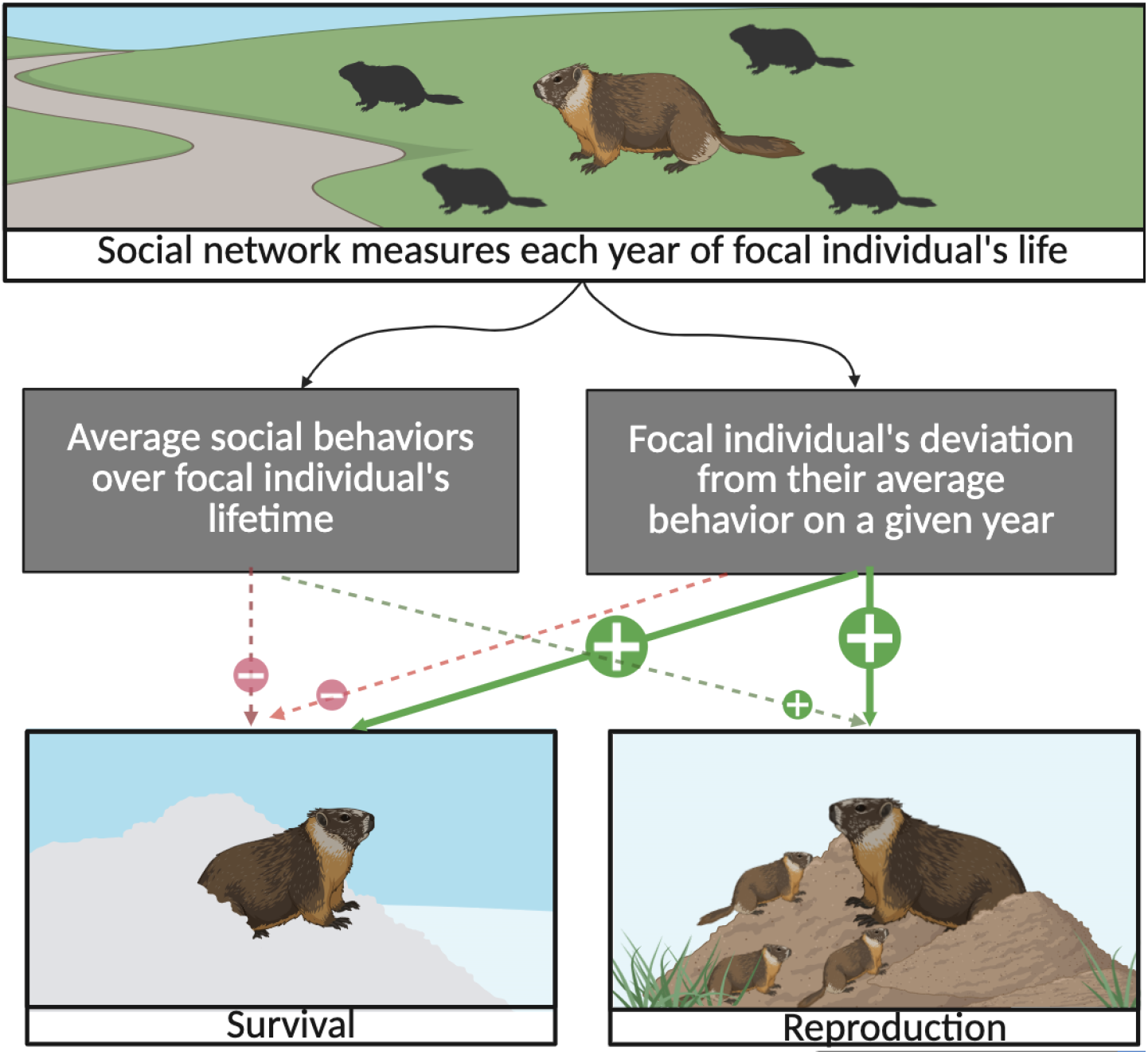

**Highlights:**

- Lifetime averages of social network traits are not associated with fitness
- Annual deviations from lifetime average sociality improves survival and reproduction
- Intra-individual behavioral plasticity may be an understudied fitness driver
- Within-individual centering is a useful approach for the study of animal behavior

## INTRODUCTION

The longevity, reciprocity, and quantity of an individual’s interactions has important fitness consequences across taxa. Social bonds are known to impact longevity and reproductive success in primates^1,2,3^, reproductive success in birds^4^ and wild horses^5^, and calving success in dolphins^6^. Across many taxa, more socially connected individuals live longer lives^3,7,8,9,10^. Yet, the individual consequences of sociality have often been studied in the context of the drivers and impacts of sociality when measured over the course of a year or season. While a valuable question, the use of annual measures may not capture the behavioral variability that may be expressed beyond a single year or season. Studies that follow individuals throughout their lifetime create additional ways to examine sociality. For example, while some individuals may maintain consistent social profiles over their lifetimes^11,12,13^, other individuals may alter their behavior over time and exhibit different social patterns across seasons and years^14,15^. Thus, in a given year, individuals may deviate from their lifetime average sociality. Here we explore whether these deviations have fitness consequences.

Recent work has identified the need to separate within-individual variation, which occurs over time in response to external and internal variables, from among-individual variation, which occurs within a population because individuals consistently differ from each other^16,17^. In the context of social relationships, within-individual effects may quantify behavioral plasticity and changes in sociality from one time period to the next. Among-individual effects quantify consistent personality and behavioral differences between individuals in a group across multiple seasons or over the lifespan of individuals (Figure 1).

**Figure 1.**
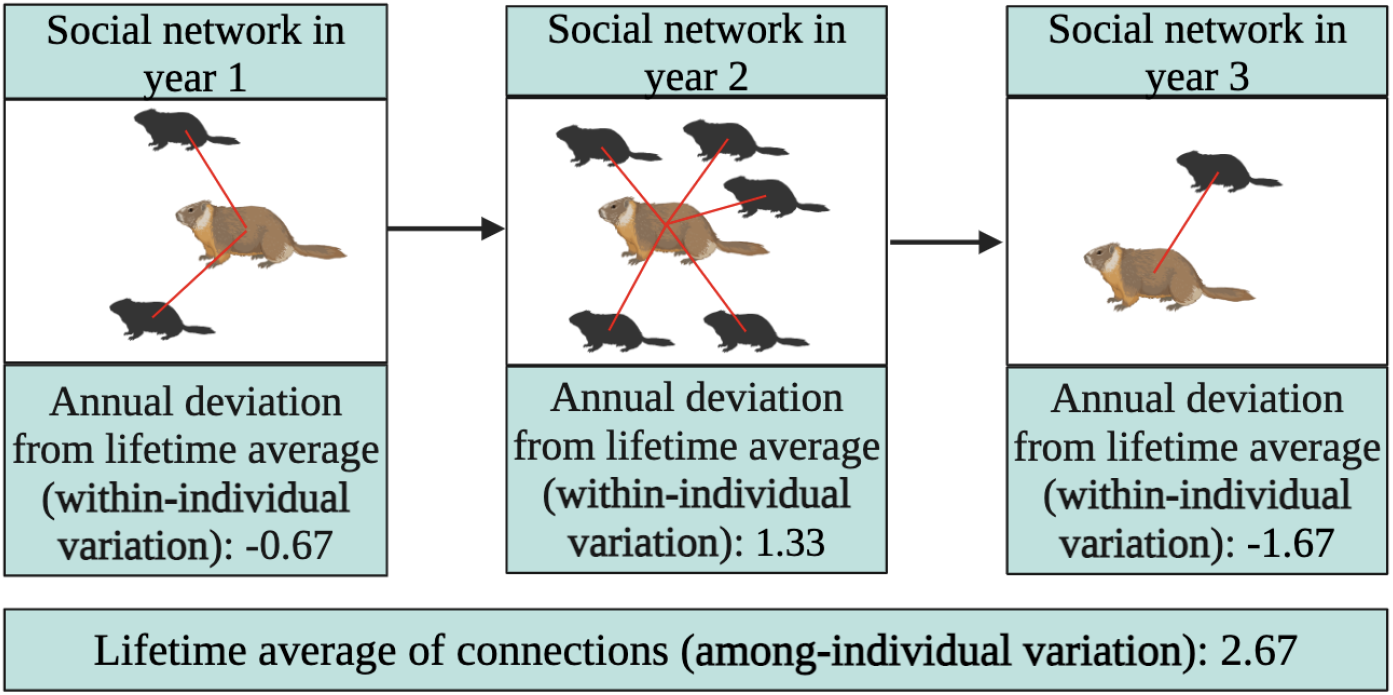
Diagram of within- and among-individual variation across an individual’s lifetime. A social network measure (in this example, number of social connections) is calculated for each year of an individual’s life. Following this, that individual’s lifetime average of connections, or among-individual variation, is calculated. Finally, that individual’s annual deviation from their lifetime average (within-individual variation) is calculated for each year by subtracting their lifetime average from their annual trait value.

Comparing within- and among-individual differences is crucial because variation at these two levels may have unique fitness consequences. For example, an individual that has consistently been very social across its lifetime, but was dramatically less social for a given season, might have a very different fitness outcome than an individual who has consistently been less social over its lifetime and continues to act asocially, despite both individuals having the same social position during that period. Persistent and stable social relationships yield benefits^3^ that may not be reflected in shorter-term, less stable relationships. Individuals that increase sociality beyond their average tendencies may also do so to reflect conditions, such as abundant resources or lack of predation, that allow a greater allocation of time and energy to social activities^18^. As such, this increase in social behavior may indicate positive conditions for both fitness and the additional fitness benefits of sociality. However, past methods of study have traditionally quantified a snapshot of an individual’s social behavior at a single point in time, which may not capture the nuance of how that individual’s lifetime behavior patterns may impact social status and fitness. The context of an individual’s deviation from their behavioral norms provides information that may impact their fitness.

Indeed, the separation of within- and among-individual differences is critical for a comprehensive understanding of other ecological and evolutionary questions^17^. Within-centering analyses have enabled advancements in our understanding of foraging effort and site fidelity^19^, movement predictability and poaching risk^20^, antipredator responses^21^, and niche specialization^22^. In the field of human behavior, individual centering methods have also been discussed as a useful method to understand personality and behavioral variation^23,24,25,26.^

Applying a within-individual centering approach to the study of social relationships in animals may lead to novel and important insights into the adaptive consequences of social behavior in wild systems. This approach is especially valuable because social behavior and relationships are often context-dependent and plastic over time^27,28^. Thus, ignoring such variation might bias results and hinder our understanding of the role of sociality and its evolution in nature.

To study the distinct processes through which relationships drive fitness over multiple seasons, we adopted a within-individual centering approach which will permit us to quantify which patterns are due to individuals modifying their behavior from previous tendencies and which patterns are due to lifetime differences among individuals within a population. To do so, we have quantified an individual’s **average social network position** over their lifetime by calculating the average network trait value across each individual’s adult years (age 2+), and then quantified each individual’s **annual deviation from this average** for 6 social network traits (Table S1). These two measures have been analyzed as separate traits which may (or may not) have fitness consequences. In addition, the annual deviations were modeled using a broken-stick approach^29^, allowing us to independently estimate the effects of positive and negative deviations from an individual lifetime average. While current work often controls for individual identity, thus somewhat accounting for inter-individual variation, the within-individual centering approach allows for intra-individual behavioral plasticity to be analyzed as a potential fitness driver.

We used this approach to evaluate the fitness consequences of both within- and among-individual variation in a population of exceptionally well-studied yellow-bellied marmots *(Marmota flaviventer*). The system is particularly well-suited to an individual centering approach because of its decades-long database^30^, its high variation in sociality both among individuals and across years, and its well-supported knowledge of the drivers of fitness. Prior work in this system has shown that individual social network position varies as a function of snowmelt and the length of the growing season^31^, which creates some degree of annual variation in individuals.

Additionally, social network position is associated with variation in yearling dispersal^32^, flight initiation distance^33^, survival^34^, longevity^35^, and reproduction^36^. However, prior work has not used a within-individual centering approach.

Given the facultatively social nature of marmots^30^ and the context-dependent benefits of sociality previously found in this system, we hypothesized that an individual’s deviation from average behavior patterns and tendency to adjust social relationships based on their surroundings (within-individual variation) will be more closely associated with increases in fitness than its average interactions or relationships (among-individual variation). This is because the benefits of sociality may outweigh its costs in many conditions. For example, in high-predation years, social relationships may yield large benefits that make sociality a good investment of time and energy^36^. In other conditions, such as years with high-density populations or short foraging seasons, high amounts of social engagement may increase competition or decrease ability to gain mass before hibernation, thus becoming maladaptive^37,38^. Condition-dependent consequences of sociality may mean that maintaining consistent social relationships across an individual’s lifetime could be costly, while deviating from an individual’s average sociality may reflect adaptive plasticity in behavior that allows individuals to modify sociality to adaptively match environmental and social contexts. Indeed, variation in the environment selects for plasticity^39^. This means that an individual’s deviation from average behaviors may be more likely to be associated with positive fitness outcomes than the average behaviors themselves.

## RESULTS

While results varied between specific social network measures, we generally found that deviations from average sociality increased the odds of survival and altered the odds of reproduction. Deviations from average sociality were associated with increased number of weaned offspring in males, where increased lifetime average sociality was associated with increased number of offspring in females.

### Within-individual deviations from lifetime average social traits were associated with increased survival in females

Deviations from an individual’s lifetime average social network position, both positive and negative, were associated with increased summer survival in females for all 6 network traits; decreases from lifetime average closeness centrality were the only deviations not associated with summer survival (Table 1, S2). Similarly, within-individual increases in all network traits and decreases in all network traits but closeness centrality were associated with increased winter survival (Table 1, S3).

**Table 1.** Abbreviated results of survival models. Only the effects of social network measures are included; unabbreviated results can be found in tables S2 and S3. Values in bold indicate statistical significance (P < 0.05). It is important to note that the broken-stick method we used creates results in which negative effect sizes of decreases in sociality signify positive relationships between that deviation and the response variable (see Figure 2 and S1).

|  | Summer survival |  |  | Winter Survival |  |  |
| --- | --- | --- | --- | --- | --- | --- |
|  | Increases in network trait | Decreases in network trait | Average of network trait | Increases in network trait | Decreases in network trait | Average of network trait |
| Degree (Female) | Est = 0.918; **P = 0.002** | Est = 0.918; **P = 0.041** | Est = 0.073; P = 0.207 | Est = 0.591; **P = 0.0002** | Est = -0.292; **P = 0.005** | Est = -0.004; P = 0.943 |
| Strength (Female) | Est = 0.049; **P = 0.015** | Est = -0.031; **P = 0.043** | Est = 0.004; P = 0.449 | Est = 0.047; **P = 0.001** | Est = -0.022; **P = 0.038** | Est = -0.008; P = 0.084 |
| Closeness centrality (Female) | Est = 34.275; **P = 0.009** | Est = -20.270; P = 0.054 | Est = 0.629; P = 0.885 | Est = 15.503; **P = 0.033** | Est = -10.477; P = 0.060 | Est = -8.137; **P = 0.049** |
| Betweenness centrality (Female) | Est = 0.072; **P = 0.011** | Est = -0.058; **P = 0.014** | Est = -0.005; P = 0.661 | Est = 0.089; **P = 0.0006** | Est = -0.070; **P = 0.001** | Est = -0.013; P = 0.227 |
| Eigenvector centrality (Female) | Est = 0.020; **P = 0.008** | Est = -0.052; **P = 0.0001** | Est = -0.002; P = 0.615 | Est = 0.022; **P = 0.003** | Est = -0.029; **P = 0.0001** | Est = -0.002; P = 0.503 |
| Clustering coefficient (Female) | Est = 5.052; **P = 0.016** | Est = -5.652; **P = 0.019** | Est = 1.050; P = 0.146 | Est = 7.572; **P = 6.35e-05** | Est = -8.288; **P = 0.0004** | Est = 0.908; P = 0.157 |
| Degree (Male) | Est = 1.741; P = 0.064 | Est = -1.646; P = 0.054 | Est = 0.132; P = 0.120 | Est = 0.576; P = 0.078 | Est = -0.500; P = 0.076 | Est = 0.034; P = 0.682 |
| Strength (Male) | Est = 0.142; P = 0.078 | Est = -0.184; **P = 0.040** | Est = -0.0008; P = 0.836 | Est = 0.082; P = 0.037 | Est = -0.080; P = 0.056 | Est = -0.021; P = 0.100 |
| Closeness centrality (Male) | Est = 5.203e+03; P = 0.998 | Est = -0.696; **P = 0.045** | Est = -1.495; P = 0.887 | Est = 18.958; P = 0.361 | Est = -56.438; P = 0.104 | Est = 3.157; P = 0.807 |
| Betweenness centrality (Male) | Est = 0.121; P = 0.059 | Est = -0.153; **P = 0.048** | Est = 0.012; P = 0.461 | Est = 0.060; P = 0.209 | Est = -0.143; **P = 0.047** | Est = 0.012; P = 0.520 |
| Eigenvector centrality (Male) | Est = 0.003; P = 0.746 | Est = -0.023; P = 0.221 | Est = -0.023; P = 0.445 | Est = 0.019; P = 0.168 | Est = -0.042; **P = 0.032** | Est = -0.006; P = 0.179 |
| Clustering coefficient (Male) | Est = 2.647; P = 0.421 | Est = -2.036; P = 0.520 | Est = -1.903; P = 0.185 | Est = 12.404; P = 0.107 | Est = -4.713; P = 0.321 | Est = -1.892; P = 0.146 |

### Within-individual deviations from lifetime average social traits were also associated with increased survival in males

Increases in male summer survival was associated with within-individual decreases in strength, closeness centrality, and betweenness centrality (Table 1, S2). Positive deviations from average strength and decreases from average betweenness centrality and eigenvector centrality were associated with winter survival in males (Table 1, S3).

### Among-individual variation had little to no association with survival

Lifetime averages of social network traits were not associated with winter or summer survival in males (Table 1). Lifetime average of closeness centrality was associated with decreased winter survival in females, but no other measure of among-individual social variation was associated with variation in female survival (Table 1, S2 and S3).

### The odds of reproduction were associated with within-individual variation in females and among-individual variation in males

Males with higher lifetime averages of clustering coefficient were more likely to reproduce on any given year (Table 2, S4). Interestingly, increases from lifetime average of clustering coefficient decreased female odds of reproduction. However, female reproduction was positively associated with positive deviations from lifetime average of degree (Table S4).

**Table 2.**
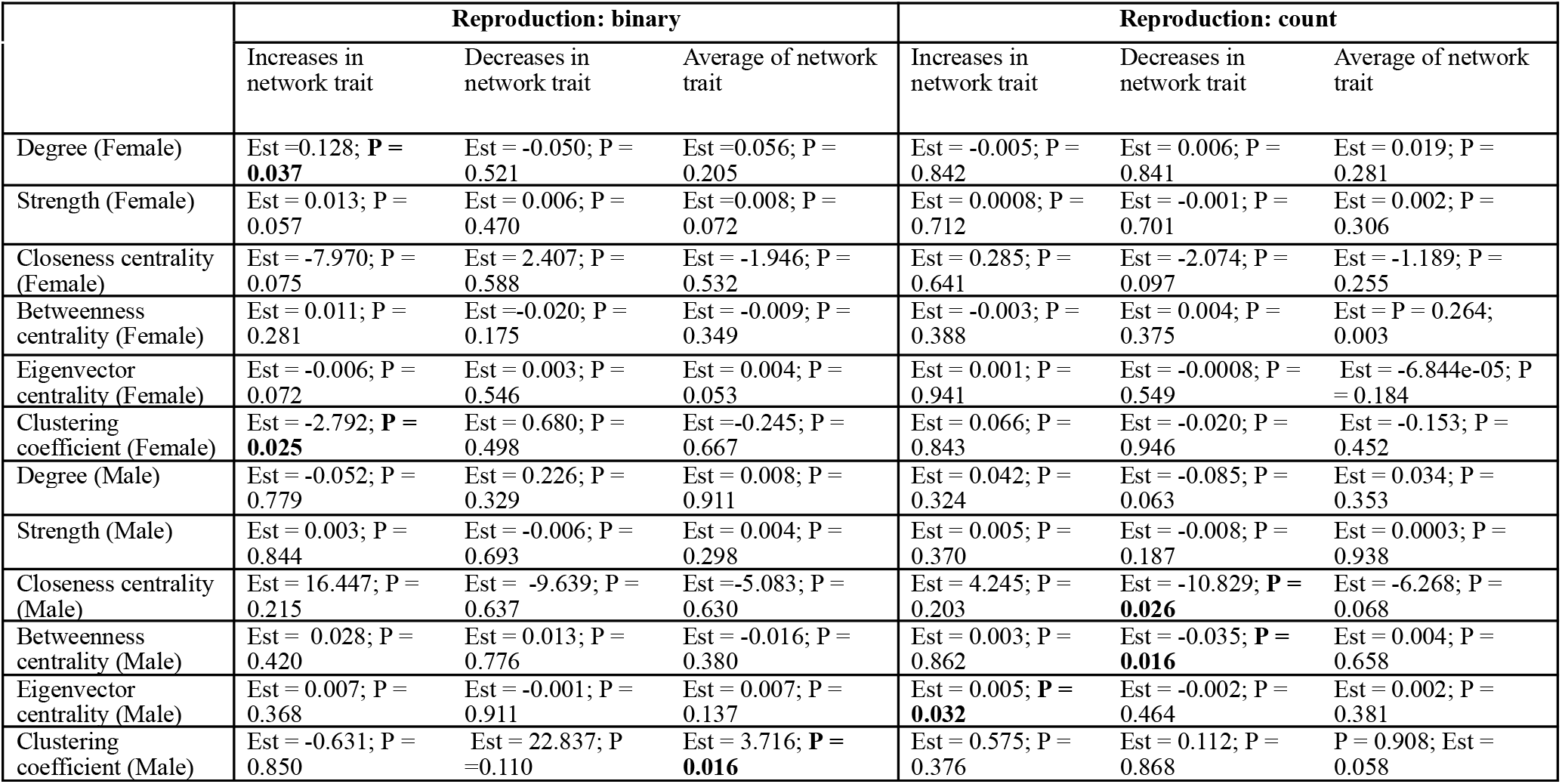
Abbreviated results of reproduction models. Only the effects of social network measures are included; unabbreviated results can be found in tables S4 and S5. Values in bold indicate statistical significance (P < 0.05). It is important to note that the broken-stick method we used creates results in which negative effect sizes of decreases in sociality signify positive relationships between that deviation and the response variable (see Figure S2 and S3).

|  | Reproduction: binary |  |  | Reproduction: count |  |  |
| --- | --- | --- | --- | --- | --- | --- |
|  | Increases in network trait | Decreases in network trait | Average of network trait | Increases in network trait | Decreases in network trait | Average of network trait |
| Degree (Female) | Est =0.128; **P = 0.037** | Est = -0.050; P = 0.521 | Est =0.056; P = 0.205 | Est = -0.005; P = 0.842 | Est = 0.006; P = 0.841 | Est = 0.019; P = 0.281 |
| Strength (Female) | Est = 0.013; P = 0.057 | Est = 0.006; P = 0.470 | Est =0.008; P = 0.072 | Est = 0.0008; P = 0.712 | Est = -0.001; P = 0.701 | Est = 0.002; P = 0.306 |
| Closeness centrality (Female) | Est = -7.970; P = 0.075 | Est = 2.407; P = 0.588 | Est = -1.946; P = 0.532 | Est = 0.285; P = 0.641 | Est = -2.074; P = 0.097 | Est = -1.189; P = 0.255 |
| Betweenness centrality (Female) | Est = 0.011; P = 0.281 | Est =-0.020; P = 0.175 | Est = -0.009; P = 0.349 | Est = -0.003; P = 0.388 | Est = 0.004; P = 0.375 | Est = P = 0.264; 0.003 |
| Eigenvector centrality (Female) | Est = -0.006; P = 0.072 | Est = 0.003; P = 0.546 | Est = 0.004; P = 0.053 | Est = 0.001; P = 0.941 | Est = -0.0008; P = 0.549 | Est = -6.844e-05; P = 0.184 |
| Clustering coefficient (Female) | Est = -2.792; **P = 0.025** | Est = 0.680; P = 0.498 | Est =-0.245; P = 0.667 | Est = 0.066; P = 0.843 | Est = -0.020; P = 0.946 | Est = -0.153; P = 0.452 |
| Degree (Male) | Est = -0.052; P = 0.779 | Est = 0.226; P = 0.329 | Est = 0.008; P = 0.911 | Est = 0.042; P = 0.324 | Est = -0.085; P = 0.063 | Est = 0.034; P = 0.353 |
| Strength (Male) | Est = 0.003; P = 0.844 | Est = -0.006; P = 0.693 | Est = 0.004; P = 0.298 | Est = 0.005; P = 0.370 | Est = -0.008; P = 0.187 | Est = 0.0003; P = 0.938 |
| Closeness centrality (Male) | Est = 16.447; P = 0.215 | Est = -9.639; P = 0.637 | Est =-5.083; P = 0.630 | Est = 4.245; P = 0.203 | Est = -10.829; **P = 0.026** | Est = -6.268; P = 0.068 |
| Betweenness centrality (Male) | Est = 0.028; P = 0.420 | Est = 0.013; P = 0.776 | Est = -0.016; P = 0.380 | Est = 0.003; P = 0.862 | Est = -0.035; **P = 0.016** | Est = 0.004; P = 0.658 |
| Eigenvector centrality (Male) | Est = 0.007; P = 0.368 | Est = -0.001; P = 0.911 | Est = 0.007; P = 0.137 | Est = 0.005; **P = 0.032** | Est = -0.002; P = 0.464 | Est = 0.002; P = 0.381 |
| Clustering coefficient (Male) | Est = -0.631; P = 0.850 | Est = 22.837; P = 0.110 | Est = 3.716; **P = 0.016** | Est = 0.575; P = 0.376 | Est = 0.112; P = 0.868 | P = 0.908; Est = 0.058 |

### Number of offspring was associated with within-individual variation in males

In males, annual number of offspring sired was positively associated with within-individual decreases in closeness centrality and betweenness centrality; number of offspring sired was also positively associated with within-individual increases in eigenvector centrality. Within-individual variation in social traits was not associated with offspring number in females (Table 1, S5).

### Non-social variables were, as expected, also associated with some fitness outcomes

Group size was negatively associated with female winter survival in the betweenness centrality model; late season body mass (August 15 mass) was positively associated with female winter survival in the betweenness centrality model (Table S3). In many models, both male and female, age was negatively associated with odds of reproduction; group size was positively associated with male odds of reproduction in the clustering coefficient model (Table S4). Mass on June 1 was also negatively associated with odds of reproduction in most models regardless of sex; however, June 1 mass was positively associated with number of offspring in females (Table S5). Group size was negatively associated with litter size in the female degree model (Table S5).

## DISCUSSION

Taken together, our results identify a strong pattern where social variability, despite having context-dependent and context-specific effects on males and females separately, is often associated with increased fitness. In line with past literature, behavioral flexibility seems to be a crucial driver of fitness^40,41,42, 43,44,45^. These results highlight the importance of examining not only an individual’s annual sociality, or their average sociality, but also an individual’s annual or seasonal deviations from its average social network position when evaluating the fitness consequences of social relationships. Our results emphasize the importance of focusing on individual deviations because, as we report, significantly altering the nature of individual social relationships has fitness consequences.

### Within-individual variation in social relationships is associated with increased survival

While an individual’s lifetime average social relationships were not generally associated with survival, deviations from an individual’s lifetime average were associated with increased summer survival in both males and females (Figure 2). This is true for both positive and negative deviations in almost every social network trait tested. Deviations from an individual’s lifetime average of most social network traits were also associated with increased winter survival in females but not males (Figure S1). This indicates that the ability to adjust one’s social relationships based on surroundings can dramatically improve survival in adult marmots, especially for females.

**Figure 2.**
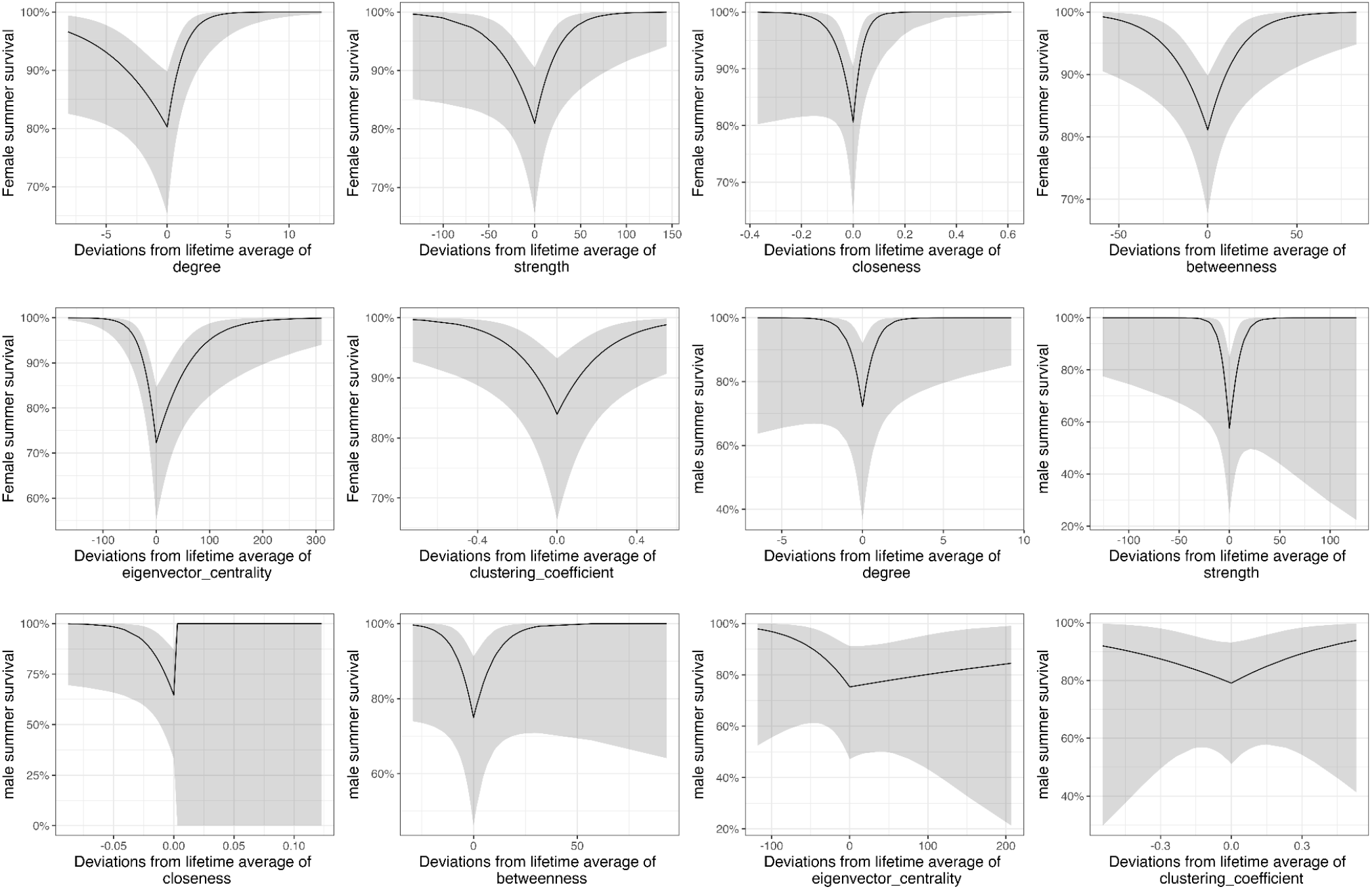
Effect of deviations in social network traits on summer survival. Deviations, both positive and negative, are associated with increased female summer survival. Decreases from lifetime averages of certain social network traits are associated with increased male summer survival.

Interestingly, these patterns are multidirectional, meaning that both increases and decreases in sociality from an individual’s lifetime average of a social network measure were associated with improved odds of survival. This indicates that the adaptive value of social relationships varies across years, and that an individual’s ability to adapt their behavior to their environmental cues is crucial in the marmot system. This flexibility seems to be much more important for survival than an individual’s inherent social tendency, at least as measured by the social network traits we quantified, because an individual’s average social network position had no association with survival.

This finding, while unique in its approach, follows past literature that has established the importance of social plasticity on individual fitness across taxa^40,41,42,43,44,45^. This pattern particularly makes sense in the marmot system, as individuals experience a variety of environmental and social conditions throughout their lifetime. Across years, an individual’s resource availability, surrounding weather patterns, predation pressure, and social group size/composition can change dramatically^46,14^. These varying social and environmental cues may alter the adaptive value of social relationships from year to year and across colony sites, causing both within-individual increases and decreases to improve survival as individuals adjust behavior to suit the cues around them.

In this system, predation is the major source of summer mortality^47^. Winter survival, on the other hand, is largely driven by gaining/maintaining body mass during the obligate 7-8 month hibernation^48^. As such, a majority of energy and time is spent on surviving the summer and gaining enough mass to survive the winter. Social relationships may have positive and negative impacts on these different needs, and our findings suggest that an individual’s ability to adjust their social network position based on their surroundings is important. For example, summer predation pressure may increase the value of social relationships, which can mitigate predation threats through detection effects or dilution effects^34,49,50^. If individuals who increase their sociality during high-predation years see improved odds of survival compared to individuals who do not adjust their behavior, this would explain why within-individual increases in social network measures are associated with increased summer survival. Females who increase their sociality from their average behaviors in response to high-predation years may be able to forage more freely than females who do not exhibit this plasticity; less-connected females who do not increase their sociality in response to predation pressure may have fewer conspecifics to warn them about predators, forcing them to spend more time vigilant and less time gaining mass to survive the winter. This could also have the consequence of social connectedness being associated with winter survival.

In the absence of high predation pressure, on the other hand, allocating time and energy to develop social relationships may impede the ability to forage^18^. Individuals able to decrease their social interactions and vigilance from their lifetime patterns in response to low-predation environments may be able to gain comparatively more mass for hibernation. It is notable that males, who are relatively less social despite being integral to social networks^51^, and do not pay the costs of reproduction, tend to gain mass much faster than females and are likely more in danger of predation than winter mortality. As such, social plasticity may be less important to males for their winter survival, explaining the lack of association between within-individual variation and winter survival in males.

### Within-individual variation in social relationships is associated with increased reproduction in males but yields mixed results in females

Males that varied from their lifetime averages of social relationships had more offspring (Figure S3). Specifically, in years where males had a higher eigenvector centrality, a lower closeness centrality, or a lower betweenness centrality than their lifetime average, they fathered more offspring. Social flexibility was not associated with the likelihood of reproducing in males (Figure S2). In contrast, within-individual variation was not associated with offspring number in females but was associated with odds of reproducing. Females were more likely to reproduce in years when they had more social partners than their lifetime average and females were less likely to reproduce in years where they had a higher clustering coefficient than average. These qualitatively different results imply that adjusting some (but not all) social patterns in line with an individual’s sex and environment may enhance reproduction. It is notable that there are fewer significant associations between within-individual social variation and reproduction than between this social variation and survival. This may imply that social plasticity is a more important driver of survival than of reproduction, especially given the mixed effects of social variability on reproduction in females.

### Within-individual variation is more strongly associated with annual fitness than lifetime patterns of behavior

An individual’s lifetime average of the vast majority of social network traits was not associated with survival or reproduction. Exceptions include those females with higher lifetime averages of closeness centrality (relative to other females) had lower annual odds of winter survival, whereas males with higher lifetime averages of clustering coefficient were more likely to reproduce each year. While this may indicate that there are some sex-specific consequences of being consistently, highly social, the lack of other significant relationships suggest that social plasticity is a more important determinant of fitness in this system. This is a unique finding that requires further study.

Because marmots are facultatively social, social plasticity may be particularly important in this system; their survival may depend more on an individual’s ability to modulate their sociality given internal and external conditions and adjust accordingly. It is possible that more socially stable, obligately social species show stronger selection for high, stable levels of sociality that remain unchanged throughout an individual’s lifetime^3,4^. However, the implications of this study reveal that the importance of social plasticity is not yet fully understood and requires further study. Indeed, this approach may be of particular interest to the study of long-lived obligately social species; social bonds are vital to the wellbeing of individuals and groups in these species, but surprisingly, social relationships (as typically quantified and studied) are frequently not found to be associated with fitness outcomes^52^. Results from within-centering approaches may change our conclusions.

### A within-individual centering approach provides key insights into the consequences of sociality

These results allow us to better understand the mechanisms behind past findings in this system and expand upon ideas presented in other study systems. Prior marmot work, which used smaller data sets and did not take an individual-centered approach, found that increased annual sociality was linked to improved female summer survival^34^, reduced overwinter survival^53^, and lower likelihood of reproduction in females^36^. These findings do not necessarily conflict with our current findings, as our analytical framework parses out lifetime patterns of sociality from annual behaviors. While being highly social in a given year may be associated with altered survival and reproduction, this association seems to be mediated through social plasticity, which has not been explicitly modelled in previous studies.

In this and other species, there may be consequences of within-individual social variation that are not detected when examining an individual’s average sociality. As such, formally quantifying intra-individual social plasticity may create new insights about the mechanisms of plasticity. Some phenotypes, especially behavioral phenotypes, may change multiple times throughout an individual’s lifetime^15^. The ability to rapidly adjust such behaviors suggests that they may have both strong fitness consequences and strong variation in the adaptive value of different phenotypes depending on environmental context. The within-individual centering approach may allow us to quantify this plasticity and study how this variation is maintained. This approach may be particularly interesting when studying obligately social species where individuals often behave consistently^12,54,55,56^ and stable, long-lasting relationships have sizable fitness benefits^3,4^. In such systems, we must also understand the fitness consequences of within-individual variation.

Additionally, a within-individual centering approach may help us further understand individuals’ capacity for and tendency to exhibit behavioral plasticity. Indeed, recent work shows that within-individual behavioral variation may have a genetic basis^57,58^. This suggests that there may indeed be selection on plasticity in the structure of social relationships. Such plasticity may be essential for survival in novel environments and will become more important as climates become more volatile^39,59^. While this study has assumed that within-individual plasticity is associated with physical environmental cues, this must be further investigated. Ultimately, social connectedness may vary as a function of social, physical, and internal cues. Further research is needed to properly understand the adaptive value of this plasticity across species.

## RESOURCE AVAILABILITY

### Lead contact

Requests for further information and resources should be directed to and will be fulfilled by the lead contact, Taylor Bastian.

### Materials availability

This study did not generate new unique reagents.

### Data and code availability

All social network and physiological data have been deposited at Open Science Framework and are publicly available as of the date of publication at DOI 10.17605/OSF.IO/5GBDZ. All original code has also been deposited at Open Science Framework and is publicly available at DOI 10.17605/OSF.IO/5GBDZ as of the date of publication. Any additional information required to reanalyze the data reported in this paper is available from the lead contact upon request.

## ACKNOWLEDGEMENTS

We thank the many decades of marmoteers for their dedication in collecting these data.

We also thank UCLA, the American Society of Mammalogists, Animal Behavior Society, American Philosophical Society, Rocky Mountain Biological Laboratory, the Natural Sciences and Engineering Research Council of Canada, the University of Ottawa, National Geographic Society, and the U.S. National Science Foundation (NSF IDBR-0754247 and DEB-1119660 and 1557130 to D.T.B.; DBI 0242960, 07211346, 1226713, and 1755522 to RMBL) for support.

Finally, we’d like to thank Greg Grether, Peter Nonacs, and Zachary Steinert-Threlkeld for their feedback on early drafts of this work.

## AUTHOR CONTRIBUTIONS

J.G.A.M and T.N.B conceived the idea. D.T.B. and J.G.A.M. supervised the project. All authors collected data. T.N.B collated data, analyzed and interpreted data and wrote the original draft with contributions from D.T.B. C.S.P contributed code necessary for creating social networks and the following social network measures. J.G.A.M assisted with statistical analysis. All authors contributed to manuscript editing.

## STAR + METHODS

### Experimental model and study participants

This study was purely observational and included the study of wild, free-living yellow-bellied marmots (*Marmota flaviventer*) in the Upper East River valley near Gunnison, CO, USA. All individuals were observed in their natural habitat and went unmanipulated. Capture and handling for sample collection occurred with permits and approval from IACUC and the Colorado Parks and Wildlife department.

This study did not involve human participants.

### Method details

Yellow-bellied marmots are facultatively social, harem-polygynous sciurid rodents^60,61^. Marmots are active for approximately five months out of the year (mid-April to mid-September) and hibernate through winter and much of spring^61^. Adult marmots reproduce soon after emerging from hibernation. Following snowmelt, and in the following summer months, they forage and gain mass in preparation for the next hibernation period^61,62^. Because body mass is associated with winter survival and emergence, mass gain is crucial during this time^63^. However, marmots experience predation pressure during months they are active above ground (Van Vuren 2001), making both foraging and socializing risky endeavors.

The marmots in and around the Rocky Mountain Biological Laboratory (38°57’N, 106°59’W; ca. 2900 m elevation) have been live-trapped and continuously studied since 1962^30,63,64,65^. During active months, trained teams of researchers ensure that virtually all individuals within our 11 colony sites are identified and regularly live-trapped. During biweekly trapping sessions, subjects are marked or re-marked, weighed, and checked for reproductive status; blood and fur samples are also taken to determine parentage and to create genealogies. Body mass measurements from each trapping event are used to calculate best linear unbiased predictions (BLUPs) by fitting linear mixed effect models to predict each individuals’ mass on June 1 and August 15^66,67^. Since 2002, detailed social observations have also been taken; during the marmots’ active hours, trained observers record all behaviors from a distance of 50-150 m through spotting scopes. The initiator, recipient, type, time, and location of all interactions are recorded. Only affiliative interactions between resident marmots (individuals recorded >5 times in a given area) are included in analyses.

### Quantification and statistical analysis

#### Quantifying Social Networks

Social network measures were calculated for each individual using interactions data. Social groups were defined through space use overlap by using SOCPROG^68^ (version 2.9) to calculate simple-ratio pairwise association indices. We then used the random walk algorithm Map Equation^69,70^ to identify social groups from these indices. Social groups included adults and yearlings, but not pups as pups typically interact only with their mothers and are difficult to observe. Because adult males may travel to matrilines outside of their primary social group to mate, adult males were included in any social groups in which they had at least one interaction. However, male social network measures were calculated within their primary group as assigned by Map Equation.

With the affiliative interactions within each group, we constructed weighted and directed networks with the R (version 4.3.3; R Core Team 2024) package “igraph”^71^ (version 2.0.2). These networks were created using affiliative interactions from mid-April till the end of June. These networks included 61,524 interactions from 538 individuals across 20 years. We calculated six social network traits, each reflecting a measure of an individual’s social relationships (Table S1).

#### Within-individual centering approach

To apply a within-individual centering approach, we quantified: 1) an individual’s lifetime average sociality using a suite of social network statistics to compare among individuals within a group, and 2) their deviation from their typical sociality to compare at the individual-level. We then fitted six sets of general or generalized linear mixed effects models. Each set compared the within-individual variation and the among-individual variation of a social network trait to one of the six isolated fitness measures (Table S1).

Social network traits were calculated for each individual for every year of their life. Because adults have very different social patterns than juveniles in this system^65,72^, we only used network measures of sexually mature individuals (i.e., those in their third year of life or later). This meant that an individual who lived for 9 years had 7 unique values of each network measure – one for every year of their adult life. Among-individual variation was quantified by calculating the average value of each social network measure over an individual’s lifetime. These values reflect their typical social behaviors and personality over their lifetime and can be compared between individuals. To quantify within-individual variation, we took the absolute value of an individual’s lifetime average of a network trait subtracted from their network value for that same trait in any given year. This measure allows each individual’s deviation from behavioral norms to be quantified and reflects an individual’s increase or decrease in sociality in a given year.

#### Statistical Analyses

We fitted generalized linear mixed models (GLMMs) in R using “lme4”^73^ for our binary response variables, winter survival and summer survival. For our reproduction model, we fitted a hurdle model in “glmmTMB”^74^ modeling the probability to reproduce and the number of offspring produced when reproducing. Due to the collinearity among social network traits, they were tested in separate models. For each social network trait, we fitted an individual lifetime average and its individual deviation from the average as fixed effects. In addition, the individual deviations were fitted with a broken-stick regression approach allowing to estimate the impact of positive and negative deviation separately. It is important to note that the broken-stick method we used creates results in which negative effect sizes of decreases in sociality signify positive relationships between that deviation and the response variable (see Figure 2 and S1). In all models, we included age, social group size, and valley location as fixed effects, all of which are known to impact social behavior and/or fitness in this system^14,34,75^. Individual ID, colony, and year were included as random effects. Other biologically relevant factors for each fitness measure were included as fixed effects in their respective models. For each colony, we determined relative predation pressure using a median split; colonies that had a higher-than-median amount of predator sightings were determined to have high predation in that year, and colonies with lower-than-median numbers of predator sightings were considered low-predation for that year. This well-tested binary predator index^76,77^ was included as a fixed effect in our summer survival models to account for the known effects of variation in predation pressure during this season^34^. Because body mass at the end of the foraging season is known to impact hibernation success and survival through the winter^78,79,80^, we included August 15 body mass as a fixed effect in our winter survival models^79,80^. Similarly, the ability to reproduce is impacted by early summer mass in this system^81^, and thus mass on June 1 was included as a fixed effect in all reproduction models. All models were fitted separately for females and males, because males follow a different life history trajectory and have different social strategies than the more settled and stationary females^72^.

Mass on June 1 and August 1 were standardized using the “scale” function in base R to allow for comparisons across models. VIF (VIF<3) was assessed with the “vif” function in the ‘car’ package^82^ (version 3.1-2). Residual diagnostics assessed through the R package “DHARMa”^83^ (version 0.4.6) indicated no major violations of model assumptions. We then visualized each significant relationship using “ggplot2”^84^ (version 3.5.0). When significant relationships between sociality and fitness were found, odds ratios and Wald confidence intervals were calculated using the “exp” and “confint” functions in base R.

### Key Resources Table

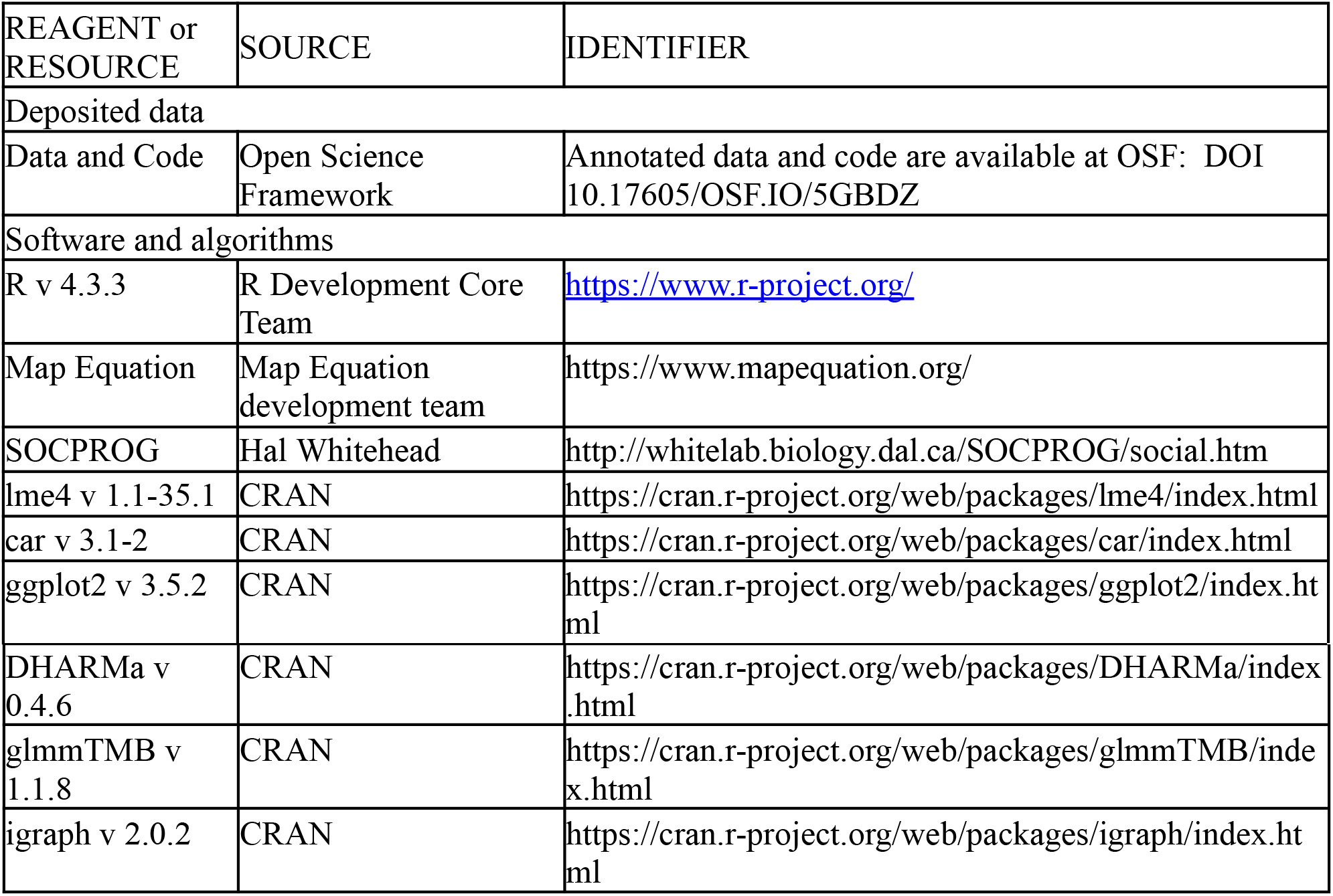

| REAGENT or RESOURCE | SOURCE | IDENTIFIER |
| --- | --- | --- |
| Deposited data |  |  |
| Data and Code | Open Science Framework | Annotated data and code are available at OSF: DOI 10.17605/OSF.IO/5GBDZ |
| Software and algorithms |  |  |
| R v 4.3.3 | R Development Core Team | https://www.r-project.org/ |
| Map Equation | Map Equation development team | https://www.mapequation.org/ |
| SOCPROG | Hal Whitehead | http://whitelab.biology.dal.ca/SOCPROG/social.htm |
| lme4 v 1.1-35.1 | CRAN | https://cran.r-project.org/web/packages/lme4/index.html |
| car v 3.1-2 | CRAN | https://cran.r-project.org/web/packages/car/index.html |
| ggplot2 v 3.5.2 | CRAN | https://cran.r-project.org/web/packages/ggplot2/index.html |
| DHARMa v 0.4.6 | CRAN | https://cran.r-project.org/web/packages/DHARMa/index.html |
| glmmTMB v 1.1.8 | CRAN | https://cran.r-project.org/web/packages/glmmTMB/index.html |
| igraph v 2.0.2 | CRAN | https://cran.r-project.org/web/packages/igraph/index.html |

## Supplemental Information

Document S1. Figures S1–S3 and Tables S1–S5

