## Supplemental Document S1 for "Deviations from an individual’s average sociality have fitness consequences"

### Supplemental Figures

**Figure S1:** Effects of deviations from lifetime average of social network traits on winter survival, related to Table 1.

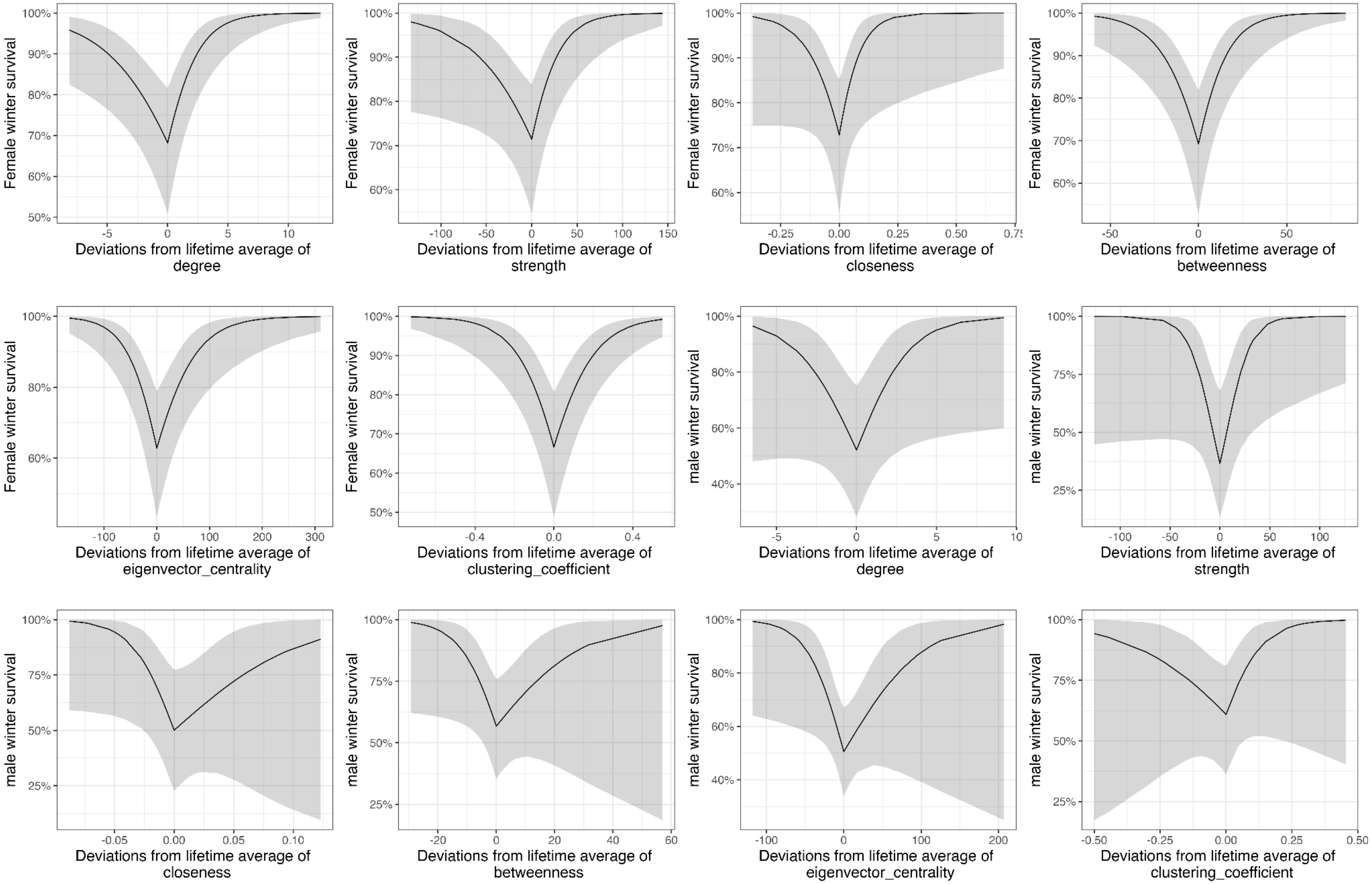

Deviations both positive and negative, are associated with increased female winter survival. Deviations from lifetime averages of social network traits are not associated with male winter survival.

**Figure S2:** Effects of deviations from lifetime average of social network traits on odds of reproducing, related to Table 2.

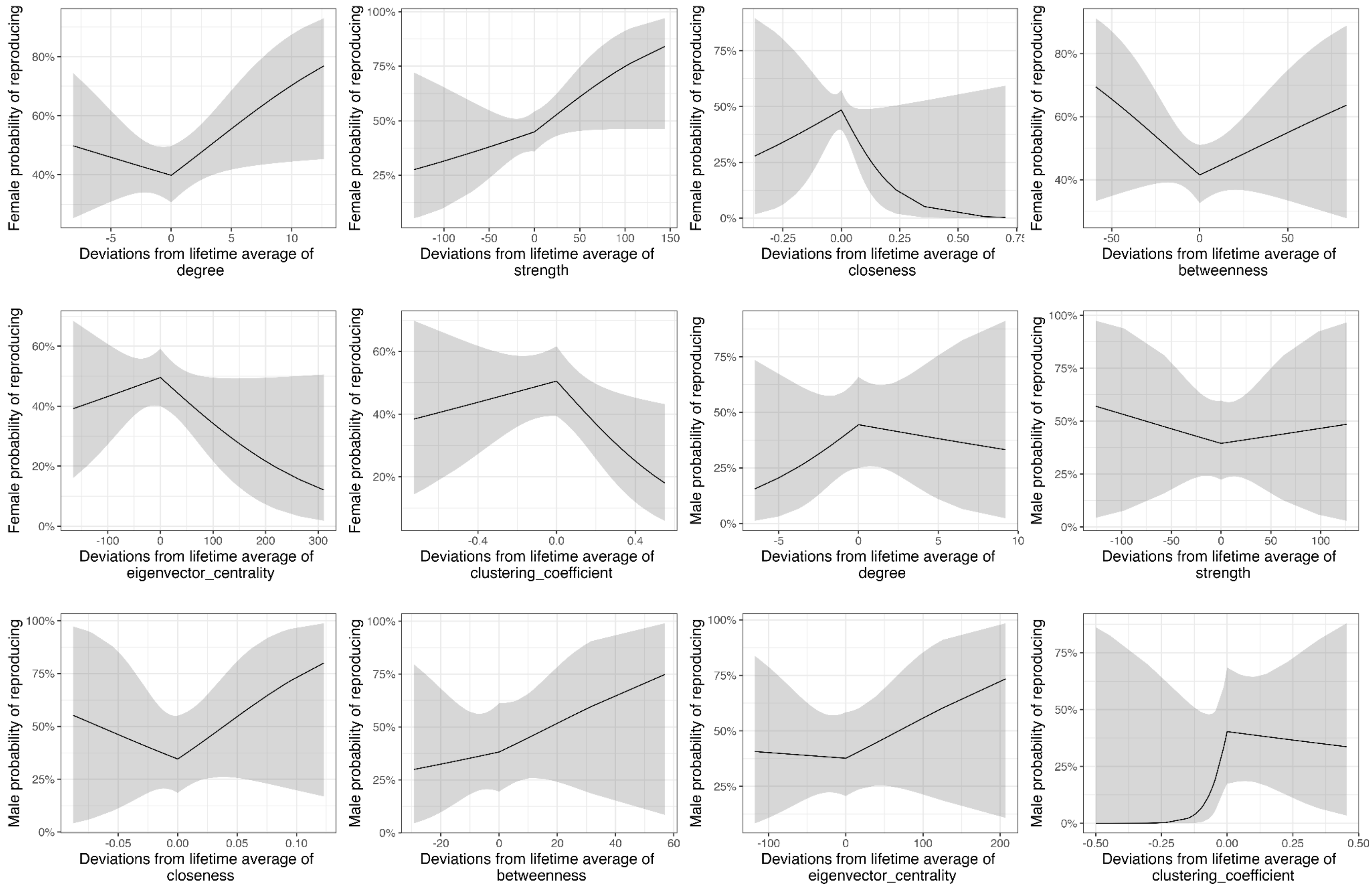

Increases from lifetime average of certain network traits are associated with increased odds of reproduction in a given year. Deviations from lifetime averages of social network traits are not associated with male odds of reproduction.

**Figure S3:** Effects of deviations from lifetime average of social network traits on number of weaned offspring in a given year, related to Table 2.

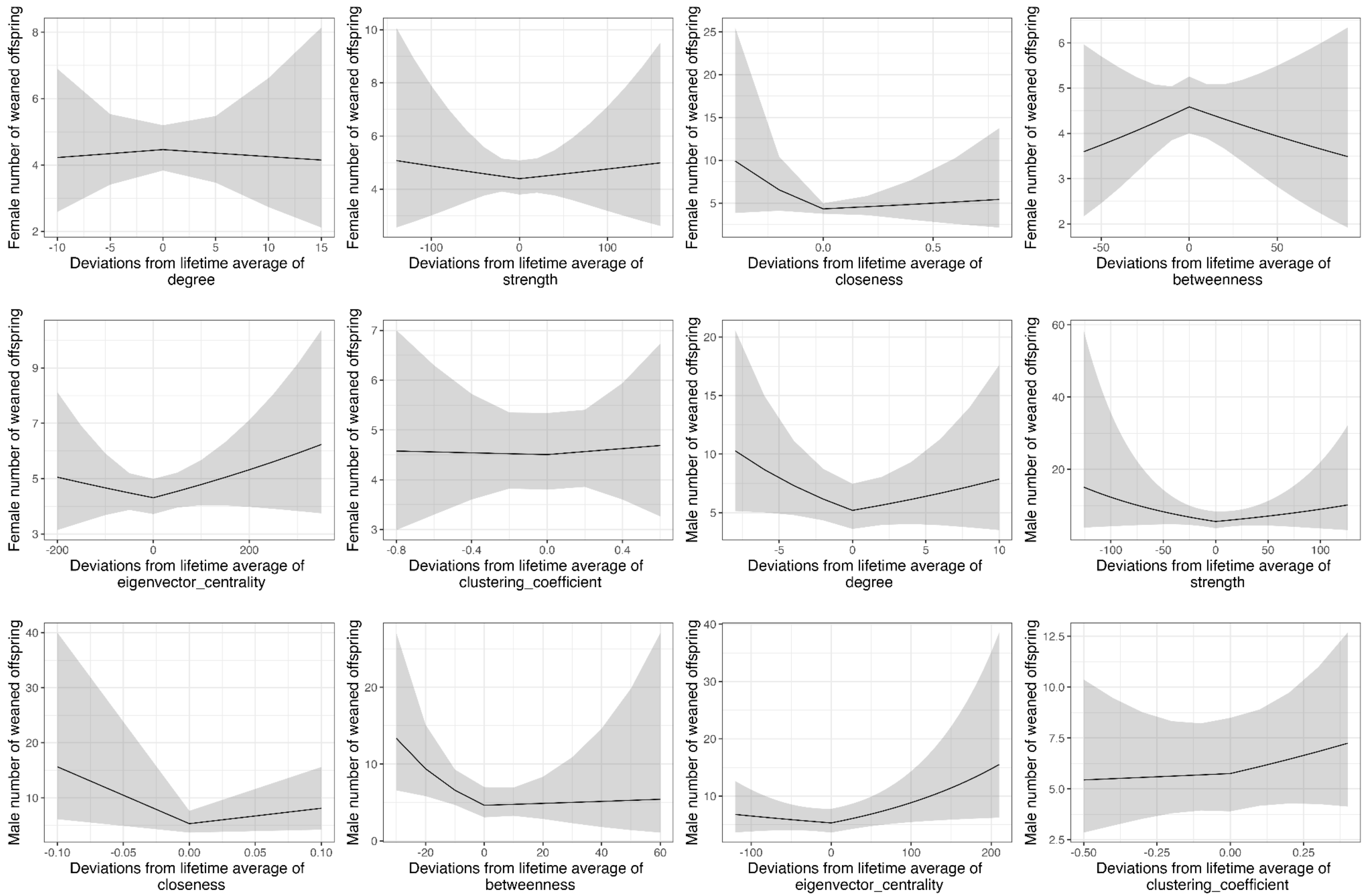

Deviations from lifetime averages of network traits are not associated with female litter size. Deviations, positive and negative, from lifetime averages of certain network traits are associated with increased number of weaned offspring in males.

**Supplemental Tables**

**Table S1**: Definitions

| Term | Description |
|---|---|
| Degree | The number of social partners a focal individual has[1] |
| Strength | The total number of interactions that a focal individual has[1] |
| Closeness Centrality | The inverse of the shortest path lengths between the focal individual and all others in its network[2,3] |
| Betweenness Centrality | A measure of how “important” an individual is in terms of group stability, social connection, and transfer of information/diseases[2,3,4] |
| Eigenvector Centrality | How well-connected a focal individual’s social partners are[5,6] |
| Clustering Coefficient | The number of existing ties between neighbors divided by the possible number of ties between neighbors[2] |
| Summer Survival | Whether or not an individual is observed/trapped after August 1 of a given year[7] |
| Winter Survival | Whether or not an individual who was seen/trapped in August or September of a given year is seen/trapped after April 1 of the following year[8] |
| Annual Reproduction | Whether or not an individual has a weaned litter in a given year[9]; parentage is determined through daily behavioral observations, adult reproductive status during their active season, trapping of all pups, and a comprehensive pedigree as determined by microsatellite markers in the DNA of pups and adults[10,11] |
| Number of offspring | Number of offspring successfully weaned, as determined by thorough observations and immediate trapping of any pups that leave their natal burrow[9] |
| Proportional Mass Gain | The amount of mass gained between June 1 and August 15 divided by mass on June 1, as determined during trapping or predicted using BLUPS[12,13] |

**Table S2:** Estimate and p-values of fixed effects on final summer survival models, related to Table 1. Values in bold indicate statistical significance ($P < 0.05$). It is important to note that the broken-stick method we used creates results in which negative effect sizes of decreases in sociality signify positive relationships between that deviation and the response variable (see Figure 2)

| | | Positive deviations in network trait | Negative deviations in network trait | Adult lifetime average of network trait | Group size | Valley location (up valley) | Local predation index (low) | Age |
|---|---|---|---|---|---|---|---|---|
| Degree (Female) | Est ± SE | 0.918 ± 0.290 | -0.239 ± 0.117 | 0.073 ± 0.058 | -0.184 ± 0.185 | -0.644 ± 0.355 | 0.056 ± 0.381 | -0.025 ± 0.070 |
| | P-value | **0.002** | **0.041** | 0.207 | 0.321 | 0.070 | 0.882 | 0.719 |
| Strength (Female) | Est ± SE | 0.049 ± 0.020 | -0.031 ± 0.015 | 0.004 ± 0.005 | 0.036 ± 0.177 | -0.454 ± 0.340 | 0.044 ± 0.391 | -0.011 ± 0.070 |
| | P-value | **0.015** | **0.043** | 0.449 | 0.840 | 0.181 | 0.910 | 0.873 |
| Closeness centrality (Female) | Est ± SE | 34.275 ± 13.040 | -20.270 ± 10.539 | 0.629 ± 4.334 | 0.334 ± 0.250 | -0.366 ± 0.412 | 0.097 ± 0.437 | -0.027 ± 0.066 |
| | P-value | **0.009** | 0.054 | 0.885 | 0.182 | 0.374 | 0.824 | 0.681 |
| Betweenness centrality (Female) | Est ± SE | 0.072 ± 0.028 | -0.058 ± 0.023 | -0.005 ± 0.011 | -0.134 ± 0.190 | -0.551 ± 0.362 | 0.13 ± 0.417 | -0.040 ± 0.068 |
| | P-value | **0.011** | **0.014** | 0.661 | 0.482 | 0.127 | 0.744 | 0.555 |
| Eigenvector centrality (Female) | Est ± SE | 0.020 ± 0.008 | -0.052 ± 0.013 | -0.002 ± 0.003 | 0.153± 0.184 | -0.263 ± 0.345 | 0.070 ± 0.380 | -0.102 ± 0.074 |
| | P-value | **0.008** | **0.0001** | 0.615 | 0.406 | 0.446 | 0.854 | 0.165 |
| Clustering coefficient (Female) | Est ± SE | 5.052 ± 2.097 | -5.652 ± 2.400 | 1.050 ± 0.721 | 0.132 ± 0.259 | -0.854 ± 0.468 | 0.422 ± 0.482 | -0.085 ± 0.095 |
| | P-value | **0.016** | **0.019** | 0.146 | 0.611 | 0.068 | 0.382 | 0.369 |

| | | | | | | | | |
|---|---|---|---|---|---|---|---|---|
| Degree (Male) | Est ± SE | 1.741 ± 0.939 | -1.646 ± 0.853 | 0.132 ± 0.085 | -0.304 ± 0.359 | -0.775 ± 0.734 | -1.003 ± 0.763 | 0.077 ± 0.159 |
| | P-value | 0.064 | 0.054 | 0.120 | 0.400 | 0.291 | 0.188 | 0.625 |
| Strength (Male) | Est ± SE | 0.142 ± 0.080 | -0.184 ± 0.090 | -0.0008 ± 0.004 | -0.284 ±0.331 | -0.101 ± 0.709 | -0.247 ± 0.688 | 0.012 ± 0.189 |
| | P-value | 0.078 | **0.040** | 0.836 | 0.392 | 0.887 | 0.719 | 0.951 |
| Closeness centrality (Male) | Est ± SE | 5.203e+03 ± 1.691e+06 | -0.696 ± 0.347 | -1.495 ± 10.500 | -0.116 ± 0.415 | -0.608 0.653 | -0.260 ± 0.619 | 0.047 ± 0.161 |
| | P-value | 0.998 | **0.045** | 0.887 | 0.781 | 0.352 | 0.674 | 0.769 |
| Betweenness centrality (Male) | Est ± SE | 0.121 ± 0.064 | -0.153 ± 0.077 | 0.012 ± 0.016 | -0.570± 0.349 | -0.764 ± 0.618 | -0.519 ± 0.619 | 0.084 ± 0.173 |
| | P-value | 0.059 | **0.048** | 0.461 | 0.102 | 0.216 | 0.401 | 0.629 |
| Eigenvector centrality (Male) | Est ± SE | 0.003 ± 0.009 | -0.023 ± 0.019 | -0.004 ± 0.006 | -0.189 ± 0.308 | -0.596 ±0.592 | 0.002 ± 0.643 | 0.279 ± 0.167 |
| | P-value | 0.746 | 0.221 | 0.445 | 0.539 | 0.313 | 0.997 | 0.095 |
| Clustering coefficient (Male) | Est ± SE | 2.647 ± 3.287 | -2.036 ± 3.161 | -1.903 ± 1.436 | -0.279 ± 0.363 | -0.468 ± 0.633 | -0.396 ± 0.622 | 0.270 ± 0.192 |
| | P-value | 0.421 | 0.520 | 0.185 | 0.443 | 0.460 | 0.525 | 0.159 |

**Table S3:** Estimate and p-values of fixed effects on final winter survival models, related to Table 1. Values in bold indicate statistical significance ($P < 0.05$). Valley location and age were removed from the male eigenvector centrality model to allow for model convergence. It is important to note that the broken-stick method we used creates results in which negative effect sizes of decreases in sociality signify positive relationships between that deviation and the response variable (see Figure S1).

| | | Positive deviations in network trait | Negative deviations in network trait | Adult lifetime average of network trait | Group size | Valley location (upvalley) | August mass | Age |
|---|---|---|---|---|---|---|---|---|
| Degree (Female) | Est ± SE | 0.591 ± 0.157 | -0.292 ± 0.105 | -0.004 ± 0.051 | -0.322 ± 0.167 | -0.392 ± 0.338 | 0.295 ± 0.165 | -0.075 ± 0.070 |
| | P-value | **0.0002** | **0.005** | 0.943 | 0.054 | 0.246 | 0.075 | 0.287 |
| Strength (Female) | Est ± SE | 0.047 ± 0.015 | -0.022 ± 0.011 | -0.008 ± 0.005 | -0.140 ± 0.157 | -0.268 ± 0.318 | 0.307 ± 0.166 | -0.055 ± 0.069 |
| | P-value | **0.001** | **0.038** | 0.084 | 0.374 | 0.399 | 0.065 | 0.428 |
| Closeness centrality (Female) | Est ± SE | 15.503 ± 7.265 | -10.477 ± 5.561 | -8.137 ± 4.129 | -0.296 ± 0.234 | -0.259 ± 0.319 | 0.265 ± 0.163 | -0.036 ± 0.068 |
| | P-value | **0.033** | 0.060 | **0.049** | 0.207 | 0.417 | 0.105 | 0.592 |
| Betweenness centrality (Female) | Est ± SE | 0.089 ± 0.026 | -0.070 ± 0.022 | -0.013 ± 0.011 | -0.355 ± 0.174 | -0.368 ± 0.321 | 0.326 ± 0.165 | -0.088 ± 0.070 |
| | P-value | **0.0006** | **0.001** | 0.227 | **0.041** | 0.252 | **0.048** | 0.206 |
| Eigenvector centrality (Female) | Est ± SE | 0.022 ± 0.007 | -0.029 ± 0.008 | -0.002 ± 0.003 | -0.057± 0.169 | -0.122 ± 0.328 | 0.224 ± 0.162 | -0.104 ± 0.072 |
| | P-value | **0.003** | **0.0001** | 0.503091 | 0.735 | 0.710 | 0.167 | 0.148 |
| Clustering coefficient (Female) | Est ± SE | 7.572 ± 1.893 | -8.288 ± 2.345 | 0.908 ± 0.642 | -0.023 ± 0.214 | -0.421 ± 0.373 | 0.239 ± 0.180 | -0.092 ± 0.077 |
| | P-value | **6.35e-05** | **0.0004** | 0.157 | 0.914 | 0.258 | 0.184 | 0.229 |

| | | | | | | | | |
|---|---|---|---|---|---|---|---|---|
| Degree (Male) | Est ± SE | 0.576 ± 0.327 | -0.500 ± 0.281 | 0.034 ± 0.082 | -0.344 ± 0.349 | 0.048 ± 0.699 | 0.865± 0.484 | -0.323 ± 0.283 |
| | P-value | 0.078 | 0.076 | 0.682 | 0.325 | 0.946 | 0.074 | 0.253 |
| Strength (Male) | Est ± SE | 0.082 ± 0.039 | -0.080 ± 0.042 | -0.021 ± 0.016 | -0.397 ± 0.321 | 0.303 ± 0.729 | 0.804 ± 0.4796 | -0.353 ± 0.268 |
| | P-value | **0.037** | 0.056 | 0.100 | 0.216 | 0.678 | 0.093 | 0.189 |
| Closeness centrality (Male) | Est ± SE | 18.958 ± 20.726 | -56.438 ± 34.751 | 3.157 ± 12.907 | -0.538 ± 0.556 | 0.487 ± 0.842 | 1.051 ± 0.541 | -0.412 ± 0.304 |
| | P-value | 0.361 | 0.104 | 0.807 | 0.333 | 0.563 | 0.052 | 0.176 |
| Betweenness centrality (Male) | Est ± SE | 0.060 ± 0.048 | -0.143 ± 0.072 | 0.012 ± 0.019 | -0.461 ± 0.324 | -0.180 ± 0.614 | 0.653 ± 0.396 | -0.151 ± 0.183 |
| | P-value | 0.209 | **0.047** | 0.520 | 0.155 | 0.770 | 0.099 | 0.409 |
| Eigenvector centrality (Male) | Est ± SE | 0.019 ± 0.014 | -0.042 ± 0.020 | -0.006 ± 0.005 | -0.195 ± 0.301 | - | 0.608 ± 0.351 | - |
| | P-value | 0.168 | **0.032** | 0.179 | 0.517 | - | 0.0832 | - |
| Clustering coefficient (Male) | Est ± SE | 12.404 ± 7.688 | -4.713 ± 4.752 | -1.892 ± 1.301 | -0.454 ± 0.412 | -0.051 ± 0.687 | 0.810 ± 0.520 | -0.074 ± 0.263 |
| | P-value | 0.107 | 0.321 | 0.146 | 0.270 | 0.940 | 0.119 | 0.779 |

**Table S4** Estimate and p-values of fixed effects on the binary half of the reproduction hurdle models, which models the probability of reproduction; related to Table 2. Values in bold indicate statistical significance ($P < 0.05$). It is important to note that the broken-stick method we used creates results in which negative effect sizes of decreases in sociality signify positive relationships between that deviation and the response variable (see Figure S2).

| | | Positive deviations in network trait | Negative deviations in network trait | Adult lifetime average of network trait | Group size | Valley location (upvalley) | June mass | Age |
|---|---|---|---|---|---|---|---|---|
| Degree (Female) | Est ± SE | 0.128 ± 0.061 | -0.050 ± 0.078 | 0.056 ± 0.044 | -0.045 ± 0.122 | -0.193 ± 0.276 | -0.481 ± 0.158 | -0.154 ± 0.066 |
| | P-value | **0.037** | 0.521 | 0.205 | 0.713 | 0.485 | **0.002** | **0.020** |
| Strength (Female) | Est ± SE | 0.013 ± 0.007 | 0.006 ± 0.008 | 0.008 ± 0.004 | 0.081 ± 0.117 | -0.031 ± 0.269 | -0.466 ± 0.161 | -0.122 ± 0.065 |
| | P-value | 0.057 | 0.470 | 0.072 | 0.487 | 0.907 | **0.004** | 0.061 |
| Closeness centrality (Female) | Est ± SE | -7.970 ± 4.478 | 2.407 ± 4.446 | -1.946 ± 3.115 | -0.142 ± 0.164 | -0.145 ± 0.267 | -0.451 ± 0.158 | -0.133 ± 0.063 |
| | P-value | 0.075 | 0.588 | 0.532 | 0.384 | 0.587 | **0.004** | **0.035** |
| Betweenness centrality (Female) | Est ± SE | 0.011 ± 0.010 | -0.020 ± 0.015 | -0.009 ± 0.009 | 0.062 ± 0.131 | -0.080 ± 0.263 | -0.503 ± 0.155 | -0.143 ± 0.065 |
| | P-value | 0.281 | 0.175 | 0.349 | 0.638 | 0.761 | **0.001** | **0.027** |
| Eigenvector centrality (Female) | Est ± SE | -0.006 ± 0.004 | 0.003 ± 0.004 | 0.004 ± 0.002 | 0.053 ± 0.121 | -0.098 ± 0.267 | -0.494 ± 0.158 | -0.010 ± 0.064 |
| | P-value | 0.072 | 0.546 | 0.053 | 0.664 | 0.714 | **0.002** | 0.118 |
| Clustering coefficient (Female) | Est ± SE | -2.792 ± 1.243 | 0.680 ± 1.004 | -0.245 ± 0.569 | -0.006 ± 0.149 | -0.255 ± 0.301 | -0.598 ± 0.179 | -0.083 ± 0.069 |
| | P-value | **0.025** | 0.498 | 0.667 | 0.966 | 0.397 | **0.0008** | 0.228 |

| | | | | | | | | |
|---|---|---|---|---|---|---|---|---|
| Degree (Male) | Est ± SE | -0.052 ± 0.184 | 0.226 ± 0.231 | 0.008 ± 0.074 | 0.314 ± 0.295 | -0.282 ± 0.567 | -0.565 ± 0.341 | -0.496 ± 0.235 |
| | P-value | 0.779 | 0.329 | 0.911 | 0.287 | 0.619 | 0.098 | **0.035** |
| Strength (Male) | Est ± SE | 0.003 ± 0.015 | -0.006 ± 0.014 | 0.004 ± 0.004 | 0.418 ± 0.274 | -0.156 ± 0.571 | -0.603 ± 0.344 | -0.571 ± 0.242 |
| | P-value | 0.844 | 0.693 | 0.298 | 0.127 | 0.785 | 0.080 | **0.018** |
| Closeness centrality (Male) | Est ± SE | 16.447 ± 13.272 | -9.639 ± 20.435 | -5.083 10.550 | 0.443± 0.373 | -0.194 ± 0.607 | -0.686 ± 0.335 | -0.602 ± 0.267 |
| | P-value | 0.215 | 0.637 | 0.630 | 0.235 | 0.750 | **0.041** | **0.024** |
| Betweenness centrality (Male) | Est ± SE | 0.028 ± 0.034 | 0.013 ± 0.044 | -0.016 ± 0.018 | 0.421 ± 0.316 | -0.232 ± 0.572 | -0.666 ± 0.348 | -0.556 ± 0.236 |
| | P-value | 0.420 | 0.776 | 0.380 | 0.174 | 0.685 | 0.056 | **0.018** |
| Eigenvector centrality (Male) | Est ± SE | 0.007 ± 0.008 | -0.001 ± 0.010 | 0.007 ± 0.005 | 0.340 ± 0.292 | -0.191 ± 0.575 | -0.789 ± 0.356 | -0.561 ± 0.234 |
| | P-value | 0.368 | 0.911 | 0.137 | 0.244 | 0.739 | **0.027** | **0.017** |
| Clustering coefficient (Male) | Est ± SE | -0.631 ± 3.342 | 22.837 ± 14.280 | 3.716 ± 1.547 | 1.126 ± 0.494 | -1.083 ± 0.780 | -0.746 ± 0.461 | -0.598 ± 0.306 |
| | P-value | 0.850 | 0.110 | **0.016** | **0.023** | 0.165 | 0.106 | 0.051 |

**Table S5:** Estimate and P-values of fixed effects on the count half of reproduction models, which explains variation in number of weaned offspring that an individual parented in a given year; related to Table 2. Values in bold indicate statistical significance ($p < 0.05$). It is important to note that the broken-stick method we used creates results in which negative effect sizes of decreases in sociality signify positive relationships between that deviation and the response variable (see Figure S3).

| | | Positive deviations in network trait | Negative deviations in network trait | Adult lifetime average of network trait | Group size | Valley location (upvalley) | June mass | Age |
|---|---|---|---|---|---|---|---|---|
| Degree (Females) | Est ± SE | -0.005 ± 0.024 | 0.006 ± 0.028 | 0.019 ± 0.017 | -0.088 ± 0.042 | -0.078 ± 0.088 | 0.182 ± 0.055 | -0.027 ± 0.018 |
| | P-value | 0.842 | 0.841 | 0.281 | **0.038** | 0.374 | **0.0008** | 0.132 |
| Strength (Females) | Est ± SE | 0.0008 ± 0.002 | -0.001 ± 0.003 | 0.002 ± 0.002 | -0.068 ± 0.041 | -0.053 ± 0.086 | 0.178 ± 0.055 | -0.026 ± 0.018 |
| | P-value | 0.712 | 0.701 | 0.306 | 0.097 | 0.534 | **0.001** | 0.163 |
| Closeness centrality (Females) | Est ± SE | 0.285 ± 0.610 | -2.074 ± 1.251 | -1.189 ± 1.043 | -0.119 ± 0.054 | -0.031 ± 0.086 | 0.177 ± 0.055 | -0.028 ± 0.018 |
| | P-value | 0.641 | 0.097 | 0.255 | **0.027** | 0.713 | **0.001** | 0.133 |
| Betweenness centrality (Females) | Est ± SE | -0.003 ± 0.004 | 0.004 ± 0.005 | 0.003 ± 0.003 | -0.087 ± 0.045 | -0.058 ± 0.084 | 0.176 ± 0.054 | -0.023 ± 0.018 |
| | P-value | 0.388 | 0.375 | 0.264 | 0.055 | 0.488 | **0.001** | 0.203 |
| Eigenvector centrality (Females) | Est ± SE | 0.001 ± 0.0008 | -0.0008 ± 0.001 | -6.844e-05 ± 0.093 | -0.073 ± 0.044 | -0.041 ± 0.084 | 0.170 ± 0.055 | -0.029 ± 0.018 |
| | P-value | 0.941 | 0.549 | 0.184 | 0.098 | 0.629 | **0.002** | 0.114 |
| Clustering coefficient (Females) | Est ± SE | 0.066 ± 0.335 | -0.020 ± 0.290 | -0.153 ± 0.204 | -0.064 0.049 | -0.009 ± 0.095 | 0.182 ± 0.061 | -0.024 ± 0.020 |
| | P-value | 0.843 | 0.946 | 0.452 | 0.192 | 0.927 | **0.003** | 0.215 |

| | | | | | | | | |
|---|---|---|---|---|---|---|---|---|
| Degree (Males) | Est ± SE | 0.042 ± 0.042 | -0.085 ± 0.046 | 0.034 ± 0.036 | -0.018 ± 0.094 | -0.280 ± 0.246 | 0.033 ± 0.112 | 0.034 ± 0.046 |
| | P-value | 0.324 | 0.063 | 0.353 | 0.847 | 0.256 | 0.769 | 0.454 |
| Strength (Males) | Est ± SE | 0.005 ± 0.005 | -0.008 ± 0.006 | 0.0003 ± 0.004 | -0.045 ± 0.081 | -0.157 ± 0.240 | 0.055 ± 0.114 | 0.014 ± 0.043 |
| | P-value | 0.370 | 0.187 | 0.938 | 0.575 | 0.514 | 0.628 | 0.739 |
| Closeness centrality (Males) | Est ± SE | 4.245 ± 3.337 | -10.829 ± 4.861 | -6.268 ± 3.433 | -0.169 ± 0.119 | -0.154 ± 0.256 | 0.103 ± 0.107 | 0.028 ± 0.044 |
| | P-value | 0.2030 | **0.026** | 0.068 | 0.156 | 0.549 | 0.339 | 0.526 |
| Betweenness centrality (Males) | Est ± SE | 0.003 ± 0.015 | -0.035 ± 0.015 | 0.004 ± 0.009 | 0.054 ± 0.095 | -0.459 ± 0.241 | 0.032 ± 0.109 | 0.023 ± 0.044 |
| | P-value | 0.862 | **0.016** | 0.658 | 0.570 | 0.057 | 0.768 | 0.600 |
| Eigenvector centrality (Males) | Est ± SE | 0.005 ± 0.002 | -0.002 ± 0.003 | 0.002 ± 0.002 | -0.042 ± 0.083 | -0.278 ± 0.250 | 0.016 ± 0.107 | 0.033 ± 0.041 |
| | P-value | **0.032** | 0.464 | 0.381 | 0.612 | 0.267 | 0.878 | 0.421 |
| Clustering coefficient (Males) | Est ± SE | 0.575 ± 0.649 | 0.112 ± 0.670 | 0.058 ± 0.503 | -0.032 ± 0.107 | -0.226 ± 0.293 | 0.046 ± 0.123 | 0.034 ± 0.039 |
| | P-value | 0.376 | 0.868 | 0.908 | 0.767 | 0.442 | 0.706 | 0.391 |
